# ACE2 protein expression within isogenic cell lines is heterogeneous and associated with distinct transcriptomes

**DOI:** 10.1101/2021.03.26.437218

**Authors:** Emily J. Sherman, Brian T. Emmer

## Abstract

The membrane protein angiotensin-converting enzyme 2 (ACE2) is a physiologic regulator of the renin-angiotensin system and the cellular receptor for the SARS-CoV-2 virus. Prior studies of ACE2 expression have primarily focused on mRNA abundance, with investigation at the protein level limited by uncertain specificity of commercial ACE2 antibodies. Here, we report our development of a sensitive and specific flow cytometry-based assay for cellular ACE2 protein abundance. Application of this approach to multiple cell lines revealed an unexpected degree of cellular heterogeneity, with detectable ACE2 protein in only a subset of cells in each isogenic population. This heterogeneity was mediated at the mRNA level by transcripts predominantly initiated from the *ACE2* proximal promoter. ACE2 expression was heritable but not fixed over multiple generations of daughter cells, with gradual drift toward the original heterogeneous background. RNA-seq profiling identified distinct transcriptomes of ACE2-expressing relative cells to non-expressing cells, with enrichment in functionally related genes and transcription factor target sets. Our findings provide a validated approach for the specific detection of ACE2 protein at the surface of single cells, support an epigenetic mechanism *ACE2* gene regulation, and identify specific pathways associated with ACE2 expression in HuH7 cells.

## INTRODUCTION

Coronavirus disease 2019 (COVID-19), caused by the SARS-CoV-2 virus, has already claimed over two million lives worldwide and remains a major threat to public health more than a year into the global pandemic^1,2^. SARS-CoV-2 establishes infection when its spike glycoprotein directly binds to its receptor, ACE2, on the surface of host cells^3^. Disruption of the spike-ACE2 interaction prevents SARS-CoV-2 infection in both cellular and animal models^4-6^, suggesting that lowering ACE2 levels may be a promising therapeutic strategy. Later in the course of infection, however, ACE2 may play a protective role, as ACE2 deficiency in mice worsens disease severity in multiple models of acute lung injury^7-11^. The mechanism for this protective effect of ACE2 is likely mediated by its physiologic function within the renin-angiotensin system. ACE2 converts angiotensin I (AngI) and II (AngII) into Ang-(1-9) and Ang-(1-7), respectively, which in turn influence vascular tone, salt and fluid balance, inflammation, cellular proliferation, and hemostasis^12,13^. Variation among individuals in ACE2 expression may contribute to the clinical heterogeneity in COVID-19 outcomes^14-16^.

Early studies of ACE2 tissue expression relied on Northern blotting or qRT-PCR of tissue homogenates and suggested moderate or high-level expression in multiple organs^17,18^. However, recent single-cell RNA sequencing (scRNA-seq) studies suggest a more restricted expression pattern, with *ACE2* transcripts often detected at low levels in only a subset of cells of a given subtype within a tissue^19-22^. Interpretation of these findings is complicated by the limited sensitivity of scRNA-seq for the reliable detection of low abundance transcripts^23^. Studies of ACE2 at the protein level have been relatively lacking, with prior reports using different antibodies often reporting conflicting results^24-27^. Likewise, *in vitro* studies of ACE2 expression have typically relied on bulk population analysis of mRNA or protein levels, leaving the distribution of ACE2 expression among individual cells within a population unknown.

We now report our development of a flow cytometry-based assay for the sensitive and specific detection of endogenous ACE2 protein at the cell surface, enabling our identification of significant cellular heterogeneity in ACE2 expression in immortalized cells. Characterization of the ACE2-expressing subset of cells was consistent with an epigenetic model of ACE2 regulation associated with distinct transcriptome profiles and differentially activated gene networks.

## RESULTS

### Development of a flow cytometry-based assay for specific detection of cellular ACE2

To identify mammalian cell lines with endogenous expression of ACE2, we first performed ACE2 immunoblotting on a panel of 7 cell lines. A band corresponding to the expected molecular weight for ACE2 was readily visualized in Caco-2, Calu-3, HepG2, HuH7, and VeroE6 cells with minimal or no ACE2 protein detected in HEK293T or HuH7.5.1 cells (Figure 1A). Our initial attempts to detect surface ACE2 staining in these cell lines by flow cytometry were limited by an equivocal signal relative to background staining for multiple ACE2 antibodies, potentially due to either low ACE2 protein abundance or an inability of the antibody to recognize extracellular ACE2 in its native conformation. To identify an antibody with optimal specificity for human ACE2 by flow cytometry, we first engineered a HEK293T cell line with stable heterologous overexpression of ACE2 for validation testing (Figure 1B). Although several commercial ACE2 antibodies tested by immunoblotting detected a specific band of the expected electrophoretic mobility (Figure S1), only 2 of 13 exhibited significantly increased staining by flow cytometry of ACE2-overexpressing HEK293T cells relative to parental cells (Figure 1C, Figure S2, Table 1).

**TABLE 1.** Human ACE2 antibodies tested in this study.

| Human ACE2 Antibody Testing |  |  |  |  |  |
| --- | --- | --- | --- | --- | --- |
| Ab # | Vendor | Catalog # | Species | FACS:<br>Overexpressed<br>ACE2 | FACS:<br>Endogenous<br>ACE2 |
| 1 | GeneTex | GTX01160 | Rabbit | - | nd |
| 2 | Abcam | 108252 | Rabbit | - | nd |
| 3 | Abcam | 272690 | Rabbit | - | nd |
| 4 | ProteinTech | 21115-1-AP | Rabbit | - | nd |
| 5 | Sigma | SAB3500978 | Rabbit | - | nd |
| 6 | Atlas | HPA000288 | Rabbit | - | nd |
| 7 | Cell Signaling | 4355S | Rabbit | - | nd |
| 8 | R&D | AF933 | Goat | + | + |
| 9 | Novus | NBP2-72117C | Mouse | - | nd |
| 10 | Santa Cruz | SC-390851 | Mouse | - | nd |
| 11 | Santa Cruz | SC-73668 | Mouse | - | nd |
| 12 | R&D | MAB933 | Mouse | - | nd |
| 13 | R&D | MAB9332 | Mouse | +++ | +++ |

**Figure 1.**
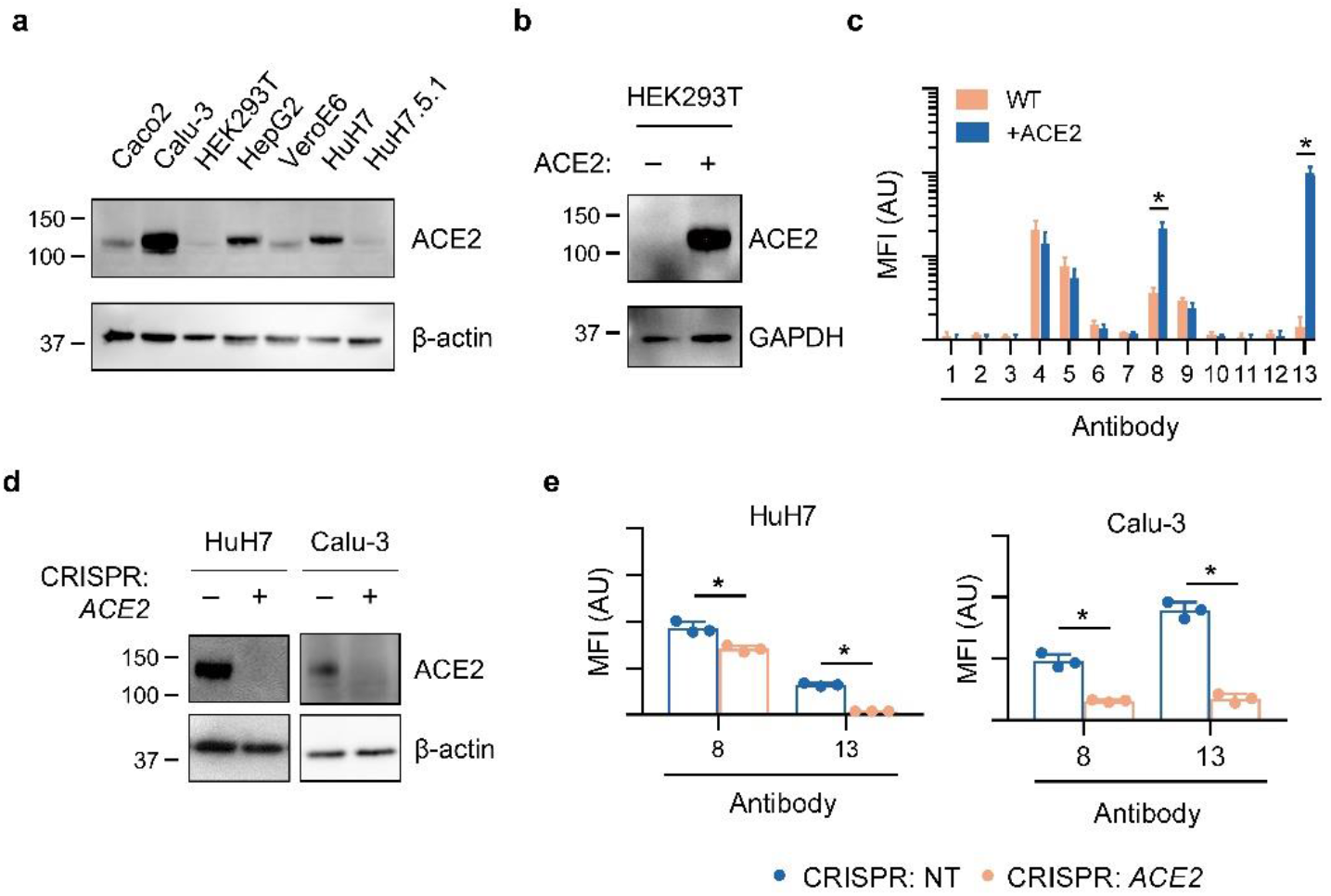
Validation of human ACE2 antibodies for flow cytometry. (A) Immunoblot of endogenous ACE2 against lysates from a panel of immortalized cell lines. (B) Immunoblot of heterologous overexpressed ACE2 in HEK293T cells. (C) Quantification of surface ACE2 mean fluorescence intensity by flow cytometry with a panel of 13 commercial antibodies against parental and ACE2-overexpressing HEK293T cells. Antibody sources are listed in Table 1. (D) Immunoblot of ACE2 in HuH7 and Calu-3 cell lines generated by CRISPR with either a nontargeting or *ACE2*-targeting gRNA. (E) Quantification of surface ACE2 mean fluorescence intensity by flow cytometry in HuH7 and Calu-3 cell lines generated by CRISPR with either a nontargeting or *ACE2*-targeting gRNA, comparing the two commercial antibodies that exhibited specific ACE2 staining in 1C. Asterisk indicates *p* < 0.05 by two-tailed Student’ s *t* test. Error bars represent standard deviation.

To assess the sensitivity of these antibodies at endogenous levels of ACE2 expression, we next applied CRISPR to generate ACE2-deficient lines of HuH7 and Calu-3 cells (Figure 1D). We confirmed that the two antibodies that had demonstrated specific staining of overexpressed ACE2 by flow cytometry were also sensitive for detection of endogenous ACE2, as indicated by increased staining of parental wild-type cells relative to ACE2-deficient lines (Figure 1E).

### Cellular ACE2 abundance is heterogeneous in multiple cell lines

Unexpectedly, in developing our flow cytometry assay, we observed a high degree of cellular heterogeneity in surface ACE2 abundance. In HuH7 cells, only ∼3-5% of cells showed increased ACE2 signal relative to unstained cells (Figure 2A). Staining in this small population was abolished in ACE2-deficient cells, indicating that this fluorescence signal was indeed specific for ACE2. Visualization of ACE2 protein in these cells by confocal microscopy supported the flow cytometry data, as we observed only a small subset of HuH7 cells with detectable surface ACE2 (Figure 2B). We observed similar heterogeneity of endogenous ACE2 surface abundance in Calu-3 cells (Figure 2C). Comparison of ACE2-positive and ACE2-negative HuH7 cells revealed no significant differences in cell size or granularity (Figure 2D) and the proportion of ACE2-expressing cells in a population was not affected by cell cycle state (Figure 2E) or cellular confluence (Figure 2F).

**Figure 2.**
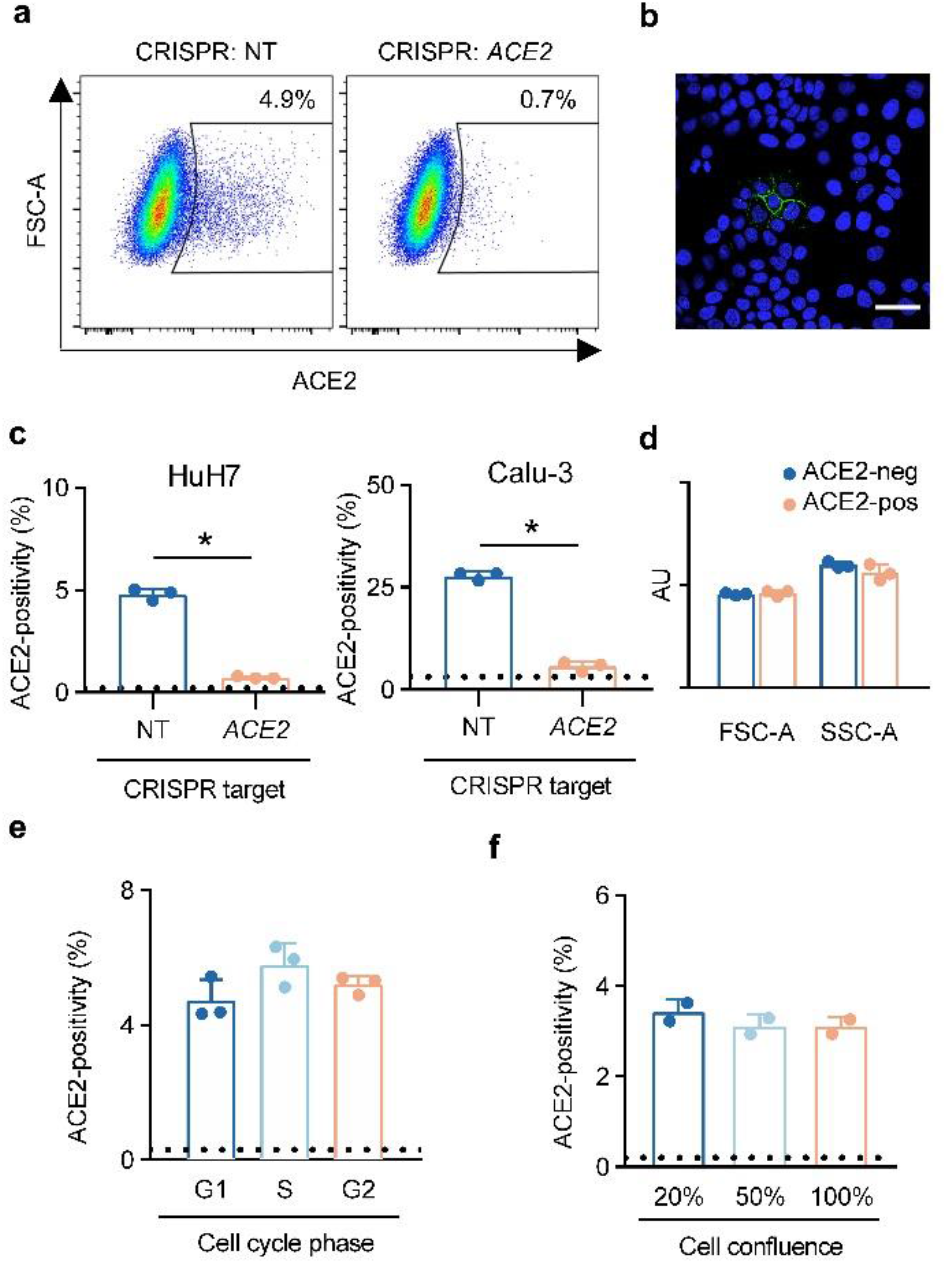
Heterogeneous ACE2 surface abundance in HuH7 and Calu-3 cells. (A) ACE2 staining by flow cytometry in HuH7 cells targeted by CRISPR with either a nontargeting or *ACE2*-targeting gRNA. (B) Confocal immunofluorescence microscopy of HuH7 cells with ACE2 (green) and Hoechst nuclear staining (blue). Scale bar = 50 µm. (C) Proportion of ACE2-positive cells as determined by flow cytometry in HuH7 and Calu-3 cells targeted by CRISPR with either a nontargeting or *ACE2*-targeting gRNA. Dashed lines represent background signal in wild-type cells stained with secondary antibody only. (D) Comparison of mean forward and side scatter parameters as determined by flow cytometry in gated ACE2-negative and ACE2-positive HuH7 cell populations. (E) ACE2-positivity by flow cytometry within each phase of the cell cycle, as determined by propidium iodide staining. (F) ACE2-positivity as determined by flow cytometry in wild-type HuH7 cells harvested at a range of cell densities. Asterisk indicates *p* < 0.05 by two-tailed Student’ s *t* test. Error bars represent standard deviation.

### HuH7 ACE2 heterogeneity is mediated at the transcript level

The cellular heterogeneity we observed for surface ACE2 protein abundance could result from differences in synthesis, turnover, or trafficking of either mRNA or protein. To distinguish among these mechanisms, we first inspected ACE2 localization in semi-permeabilized HuH7 cells by immunofluorescence. We found ACE2 protein predominantly at the plasma membrane, with no significant intracellular staining in ACE2-positive cells or in neighboring cells lacking surface ACE2 (Figure 3A). Similarly, semi-permeabilization of HuH7 cells did not significantly increase the proportion of cells with detectable ACE2 staining by flow cytometry (Figure 3B). Consistent with these findings, we also detected no ACE2 protein in lysates of sorted cells lacking surface-displayed ACE2 (Figure 3C). Analysis of mRNA from sorted cells revealed a marked reduction in *ACE2* transcripts in ACE2-negative cells (Figure 3D). Collectively, these results indicate that HuH7 cellular heterogeneity of ACE2 abundance is mediated at the mRNA level.

**Figure 3.**
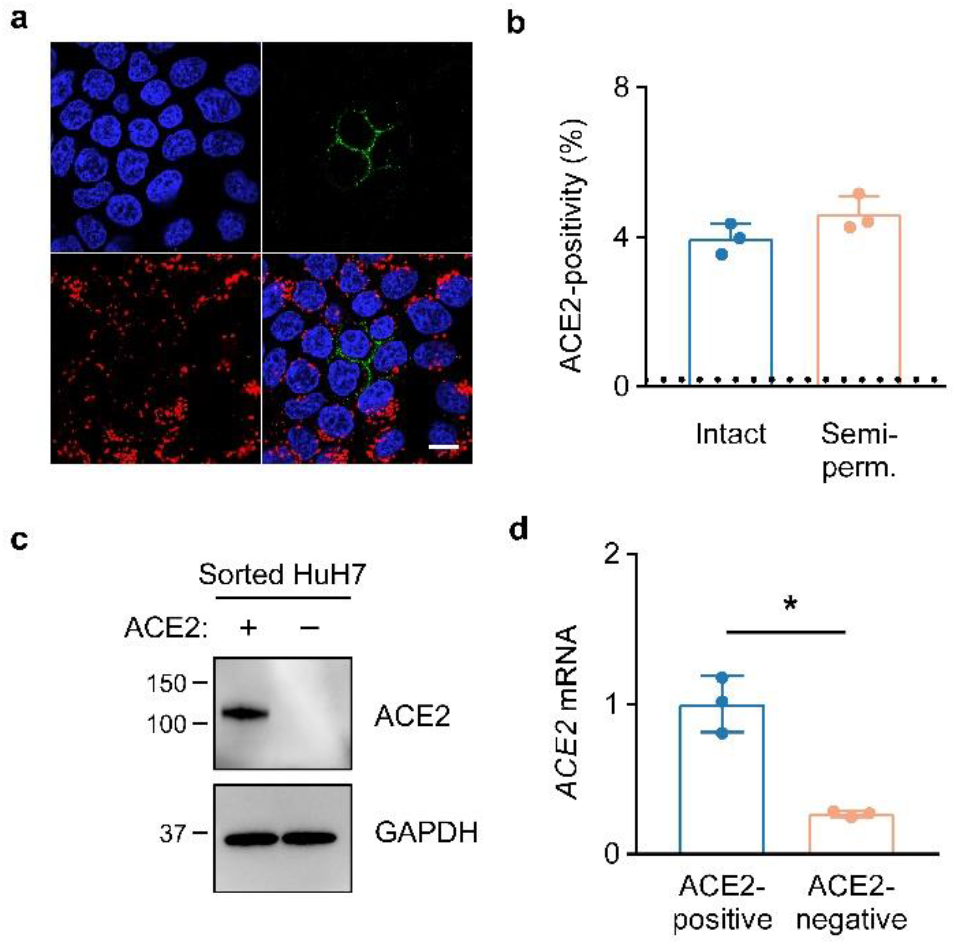
ACE2 heterogeneity in HuH7 cells is mediated at the mRNA level. (A) Immunofluorescence confocal microscopy of semi-permeabilized HuH7 cells with ACE2 (green), GM130 (red), and DAPI (blue) staining. Scale bar = 10 µm. (B) ACE2-positivity as determined by flow cytometry of intact or semi-permeabilized parental HuH7 cells. Dashed line represents background signal in cells stained only with secondary antibody. (C) Immunoblot of ACE2 in lysates from FACS-sorted ACE2-positive and ACE2-negative HuH7 cells. (D) *ACE2* transcript levels in FACS-sorted ACE2-positive and ACE2-negative HuH7 cells as determined by qRT-PCR. Asterisk indicates *p* < 0.05 by two-tailed Student’ s *t* test. Error bars represent standard deviation.

### Heritability of ACE2 expression in HuH7 cells

We noticed during immunofluorescence microscopy that ACE2-positive cells were often present in clusters (Figure 4A), suggesting that ACE2 expression in these cells may be heritable. To test this hypothesis, we sorted HuH7 cells into ACE2-positive and ACE2-negative subpopulations, expanded each in culture, and reanalyzed each population by flow cytometry. A significant difference between the sorted populations in ACE2 surface abundance was persistent after ∼7 doublings in culture (Figure 4B). ACE2-enriched cells also exhibited increased ACE2 enzymatic activity, indicating this staining to reflect functional protein (Figure 4C). Serial sorting and expansion of ACE2-positive cells led to a progressive enrichment in the proportion of ACE2-positive cells from ∼3-5% positivity in parental cells to ∼60-70% after 3 rounds of enrichment (Figure 4D). ACE2 expression, however, was not a fixed trait, as the proportion of positive cells gradually reverted back toward the parental distribution over continued passaging (Figure 4E). This decay in ACE2-positivity over time argued against selection for a somatic genetic mutation, which we confirmed by expanding 8 independent single cell clones and finding each to recapitulate the heterogeneity of the parental population, albeit at varying proportions of ACE2-positivity (Figure 4F).

**Figure 4.**
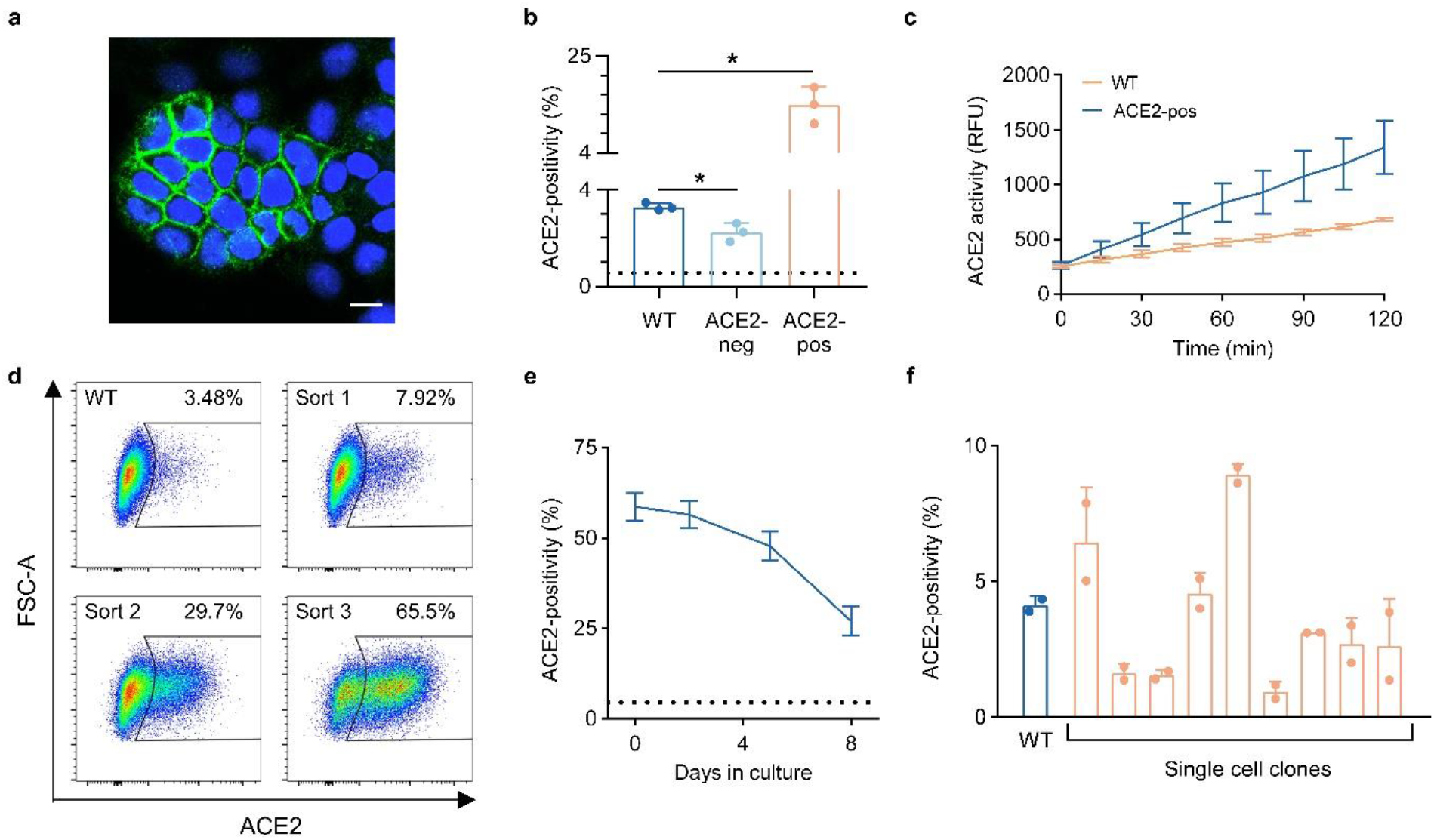
Heritability of ACE2 expression in HuH7 cells. (A) Immunofluorescence confocal microscopy showing a cluster of ACE2-positive HuH7 cells with ACE2 (green) and Hoechst nuclear staining (blue). Scale bar = 10 µm. (B) ACE2-positivity by flow cytometry in starting wild-type HuH7 cells and after ∼7 doublings in culture following FACS isolation and expansion of ACE2-positive and ACE2-negative subpopulations. Dashed line represents background signal from cells stained with only secondary antibody. (C) ACE2 enzymatic activity over 2 hours in wild-type and FACS-sorted ACE2-positive HuH7 cells. (D) Flow cytometry plots of ACE2-positivity in HuH7 cells serially enriched for ACE2 abundance over three FACS sorts. (E) Decay of ACE2-positivity at various time points of passaging after FACS enrichment of ACE2 abundance. Dashed line represents ACE2-positivity in wild-type HuH7 cells. (F) ACE2-positivity by flow cytometry in single-cell HuH7 clones compared to parental cells. Asterisk indicates *p* < 0.05 by two-tailed Student’ s *t* test. Error bars represent standard deviation.

### ACE2-expressing cells have distinct global transcriptomes relative to non-expressing cells

To investigate the molecular pathways associated with ACE2 expression, we performed RNA-seq on sorted ACE2-negative and ACE2-positive HuH7 cells. Principal component analysis demonstrated distinct and tightly clustered transcriptome profiles corresponding to independent biologic replicates for each population (Figure 5A). Differential expression analysis identified 105 genes with significantly increased (log2FC>1, *p*_adj_<0.01) transcript levels in ACE2-positive cells, and no genes with significantly decreased (log2FC<-1, *p*_adj_ <0.01) transcript levels (Figure 5B). Similar analysis of serially ACE2-enriched cells relative to parental cells also demonstrated distinct clustering profiles (Figure 5A). In comparison to the differential expression analysis between singly-sorted ACE2-negative and ACE2-positive cells, serial enrichment for ACE2 was associated with more extensive transcriptome changes, including greater numbers of genes either with increased (326) or decreased (194) transcript levels (Figure 5C). *ACE2* itself exhibited the greatest enrichment in ACE2-positive cells after a single sort, and was also among the most enriched genes after serial sorting, confirming our finding that ACE2 heterogeneity is mediated at the transcript level and validating our workflow for cell sorting and transcriptome analysis. We observed a high degree of concordance in the differentially expressed transcripts identified by either approach. Of the 105 genes identified as upregulated in the analysis of singly sorted cells, 86 were similarly identified as upregulated in serially ACE2-enriched cells (Figure 5D) with the magnitude of upregulation correlated in either approach (Figure 5E).

**Figure 5.**
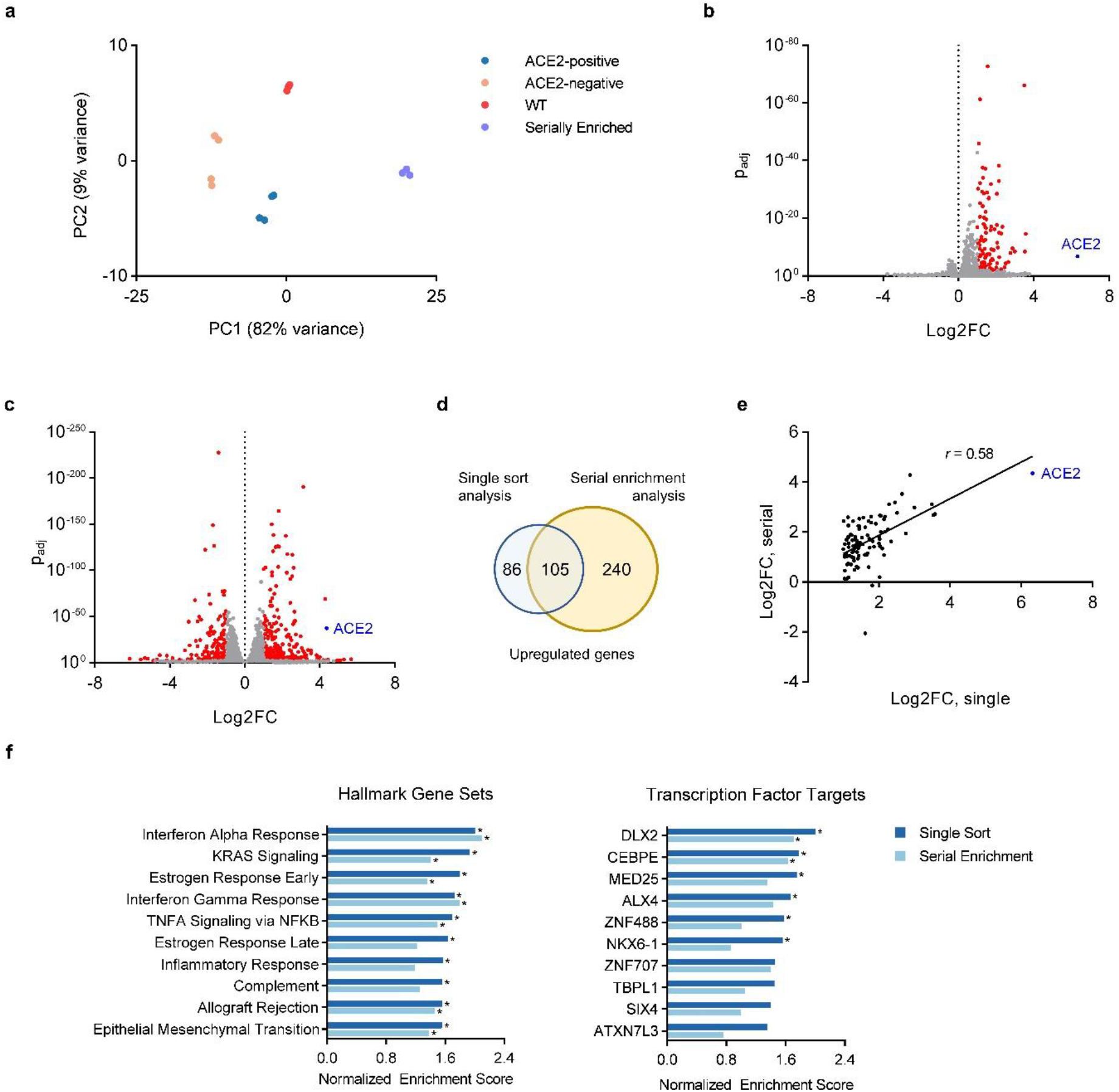
Transcriptome profiling of ACE2-expressing and non-expressing HuH7 cells. (A) Principal component analysis of RNA-seq profiles of parental wild-type cells, singly sorted ACE2-negative and ACE2-positive, and serially sorted ACE2-enriched cells. (B) Volcano plots of differentially expressed genes between ACE2-negative and ACE2-positive cells isolated after a single sort. Genes exhibiting abs(log2FC)>1 and *p*_adj_< 0.1 by DESeq2 calculation are colored in red. (C) Volcano plot of differentially expressed genes in parental HuH7 cells and serially ACE2-enriched cells. D) Venn diagram comparing the number of upregulated genes identified in each of the 2 RNA-seq analyses in 5B-C. (E) Correlation of log2 fold change for each gene upregulated in 5B in comparison to its magnitude of upregulation in 5C. (F) Gene set enrichment analysis depicting the normalized enrichment scores of the 10 most enriched hallmark pathways and transcription factors target sets in singly sorted ACE2-positive cells relative to ACE2-negative cells. The corresponding normalized enrichment score in serially sorted ACE2-enriched cells relative to wild-type cells is also plotted. Asterisk indicates FDR<25% by GSEA calculation.

### Pathway analysis of differentially expressed genes in ACE2-expressing cells

To identify transcriptome signatures correlated with ACE2 expression, we performed Gene Set Enrichment Analysis^33^ of RNA-seq data from ACE2-sorted cells. From the Molecular Signatures Database Hallmark Gene Set collection^34^, we observed significant (FDR<25%) enrichment of 17 gene sets and depletion of 2 gene sets in ACE2-positive relative to ACE2-negative cells (Figure 5F, Supplemental Table 5). Analysis of serially enriched cells relative to parental wild-type cells revealed significant enrichment for 8 hallmark gene sets, each of which had also been identified in the first analysis of singly sorted cells. Gene sets associated with ACE2 expression in both analyses included targets of interferon and estrogen signaling. Significant depletion in ACE2-enriched cells relative to wild-type cells was observed for 11 hallmark gene sets, including the 2 which had been identified after a single sort.

We also assessed for overrepresentation of transcription factor targets among ACE2-correlated genes using the Gene Transcription Regulation Database^35^. In this analysis, we found 6 transcription factor target sets with significant enrichment and none with significant depletion in ACE2-positive relative to ACE2-negative cells (Figure 5F, Supplemental Table 6). Analysis of ACE2-enriched cells identified only 2 transcription factors, DLX2 (distal-less homeobox 2) and CEBPE (CCAAT/enhancer binding protein epsilon), whose target sets were enriched in ACE2-enriched cells. DLX2 and CEBPE target gene sets were also the 2 most enriched in the analysis of singly sorted cells. No transcription factor target sets were significantly depleted in ACE2-enriched cells. Together, these findings identify candidate pathways and transcriptional regulators that are associated with ACE2 expression in HuH7 cells.

### ACE2 expression in HuH7 cells is primarily mediated by the proximal promoter for full-length transcripts

Transcription of full-length *ACE2* is initiated from either a proximal or a distal promoter with tissue-specific differences in their relative usage^28-30^. Recently, a cryptic promoter between exons 8 and 9 has also been recognized that initiates interferon-responsive transcription of a truncated, nonfunctional *ACE2* splice variant^31,32^. To clarify the relative usage of each promoter in HuH7 cells, we analyzed exon coverage among *ACE2* transcripts in our RNA-seq data of both wild-type cells and in those either singly or serially sorted based on ACE2 surface abundance. We did not observe sequences specific for the truncated *ACE2* variant for any of these samples, with no reads mapping to either exon 1c itself or the junction between exon 1c and exon 9 (Figure 6). Among full-length transcripts, we observed much fewer reads mapping to exon 1a than exon 1b and a corresponding lack of reads containing the junction of exons 1a and 1b (Figure 6). These findings suggest that *ACE2* transcription in HuH7 cells is primarily mediated by the proximal promoter of full-length splice variants.

**Figure 6.**
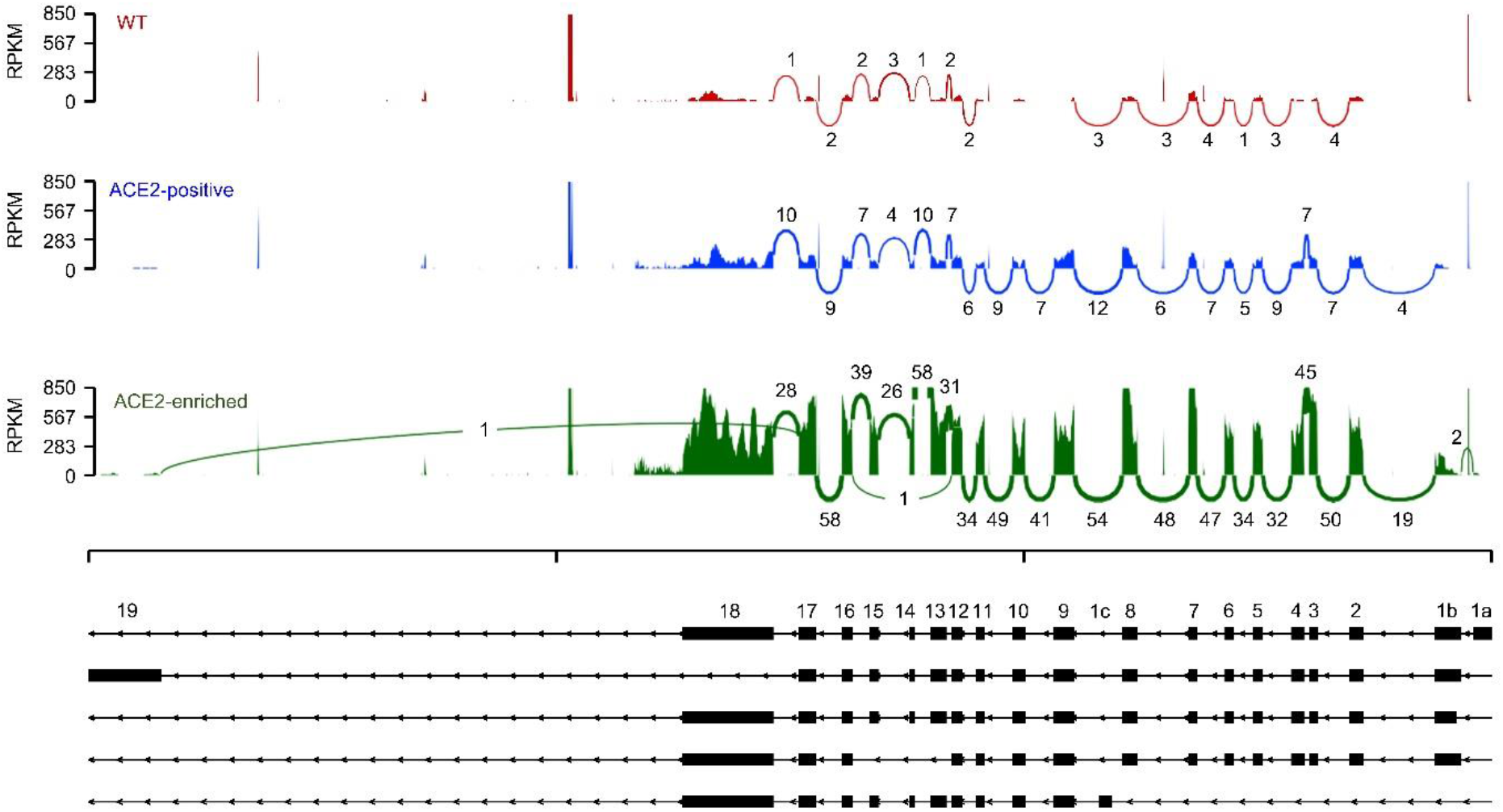
ACE2 isoform analysis in HuH7 wild-type and ACE2-enriched cells. Sashimi plot of RNA-seq mapped reads for individual *ACE2* exons and exon-exon junctions in RNA-seq of HuH7 wild-type cells, ACE2-positive cells isolated after a single sort, or serially ACE2-enriched cells. Representative plots from replicates of each sample are displayed.

## DISCUSSION

Antibodies are critical reagents for a variety of applications including immunoblotting, immunofluorescence, ELISA, and flow cytometry. Although the number of commercially available antibodies has grown dramatically, with over 5 million currently listed in the database CiteAb^36^, systematic studies have identified high failure rates, leading to serious concerns about antibody quality and calls for more stringent validation testing^37-40^.

As rigorous controls for ACE2 antibody specificity, we engineered cells overexpressing heterologous *ACE2* cDNA as well as cells with CRISPR-mediated disruption of endogenous *ACE2* gene expression. We found that 11 of 13 antibodies tested by flow cytometry demonstrated either an absence of binding above background staining, or nonspecific binding that was not influenced by ACE2 overexpression or deletion. These data do not rule out the potential of these antibodies to exhibit ACE2-specific binding in other applications, such as immunohistochemistry or immunofluorescence. Several of these antibodies indeed recognized ACE2-specific bands by immunoblotting, potentially resulting from recognition of a denatured epitope or to electrophoretic separation of cross-reactive proteins. For flow cytometry, it also remains possible that optimization of our staining protocol (e.g. antibody concentration, blocking or wash conditions) might uncover ACE2-specific binding. Under the conditions of our staining, however, only 2 antibodies tested (R&D #MAB9332 and R&D #AF933) showed clearly significant dependence of binding upon ACE2 expression. Our systematic validation of these antibodies will serve as a resource for researchers interested in either quantifying human ACE2 protein at the single cell level, or in isolating cells based upon their ACE2 protein abundance.

Our validation of ACE2-specific antibodies for flow cytometry enabled our observation of striking cellular heterogeneity in ACE2 surface abundance. This phenomenon was confirmed in multiple cell lines, suggesting physiologic relevance and consistent with scRNA-seq studies *in vivo* demonstrating heterogeneity of *ACE2* transcripts among cell types within a tissue, as well as among cells of a given subtype^19-22^. At the protein level, recent immunohistochemistry studies using strict antibody validation criteria have also detected low level and heterogeneous ACE2 protein expression in the respiratory tract and in other tissues^41,42^. Cells within a tissue have distinct developmental histories and microenvironments that may lead to broad transcriptome changes unrelated to the coincident differential expression of a given gene. Our finding of similar heterogeneity in immortalized cell lines, arising from a clonal origin and growing in the same culture dish, suggests they may serve as a simplified model to dissect the causal pathways governing ACE2 expression.

Although our findings do not define the molecular basis for ACE2 heterogeneity in HuH7 cells, a number of clues point toward an epigenetic mode of regulation. First, we found ACE2 heterogeneity to be driven at the mRNA level, since intracellular ACE2 protein was absent in cells lacking surface ACE2 while *ACE2* mRNA was reduced in these cells and progressively increased upon serial enrichment for surface ACE2 protein. Second, ACE2 expression in these cells was heritable, with daughter cells of ACE2-expressing cells maintaining expression over several generations. Third, a strictly genetic basis for this phenomenon was ruled out by the reversion of the ACE2 expression back toward the parental distribution over time, together with the recapitulation of ACE2 heterogeneity upon expansion of single cell clones. This pattern of inheritance suggests epigenetic regulation, whereby chemical modifications of chromatin govern the accessibility of the *ACE2* gene for transcription. Consistent with this model, a variety of histone and DNA modifications as well as transcription factor motifs and binding have been associated with the *ACE2* locus^43,44^.

The ability to isolate ACE2-expressing and non-expressing cells allowed us to interrogate the transcriptome profiles of each subpopulation. We identified clearly distinct transcriptomes for ACE2-expressing and non-expressing cells. Differential expression analysis identified broad transcriptome changes associated with ACE2 expression. The validity of these findings is supported by the high degree of concordance in genes identified as differentially expressed either after a single sort and upon serial ACE2 enrichment. Pathway analysis identified several different transcriptional programs associated with ACE2 expression. Among the most enriched gene sets in ACE2-expressing cells were those associated with interferon-α and interferon-γ responses. The influence of interferon signaling on ACE2 expression is controversial, with initial scRNA-seq of lung tissue suggesting an upregulation of ACE2 transcript levels by interferons^19,45^. It was subsequently discovered that a truncated, nonfunctional isoform of *ACE2* is expressed from a cryptic IFN-responsive promoter^31,32^. The ACE2 antibody we used for flow cytometry was generated against an extracellular epitope not present in the truncated isoform. Furthermore, we did not detect the truncated *ACE2* isoform by RNA-seq analysis in either wild-type or ACE2-enriched HuH7 cells. Our finding of increased expression of interferon response genes in ACE2-expressing cells was therefore not driven by an association with this truncated isoform but rather initiated from the proximal promoter of full-length isoforms.

Intriguingly, we also observed an association of ACE2 expression with both early and late responses to estrogen. Patient outcomes in COVID-19 exhibit a sexual dimorphism, with men more likely to develop severe disease and death^46^. The molecular mechanism for this observation remains uncertain, but potential regulation of ACE2 expression by estrogen is further supported by *in vitro* studies that demonstrated an ERα-dependent increase in *ACE2* mRNA by 17β-estradiol treatment in both 3T3-L1 adipocytes^47^ and HUVECs^48^. An opposite effect, however, was observed in differentiated NHBE cells, as 17β-estradiol treatment led to a decrease in *ACE2* mRNA^49^. An analysis of the Genotype-Tissue Expression (GTEx) database found a correlation across tissues between transcript levels of *ACE2* and estrogen receptors *ESR1* and *ESR2*^50^ and a bioinformatics analysis identified several ESR1 and ESR2 binding motifs in the *ACE2* promoter^30^. Our findings support a potential role of estrogen signaling in *ACE2* gene regulation.

Our analysis of transcription factors identified an association with ACE2 expression for both DLX2 and CEBPE, which represented the top 2 most enriched target sets in both independent analyses of singly sorted and serially enriched ACE2-expressing cells. DLX2 is a homeobox transcription factor that has been previously implicated in development of the forebrain, differentiation of interneurons, and regulation of TGF-β signaling^51-53^. CEBPE is a transcriptional activator with a bZIP DNA-binding domain that has been previously implicated in granulocyte differentiation and acute myeloid leukemia^54-56^. Neither DLX2 nor CEBPE has been previously associated with *ACE2* regulation, and *ACE2* itself is not among the gene sets of annotated DLX2 or CEBPE targets. It is possible that DLX2 and/or CEPBE regulate *ACE2* via a direct or an indirect mechanism. Importantly, however, a limitation of our study is that our identification of differentially expressed genes in ACE2-positive cells does not distinguish between causal and correlative relationships. Further investigation will be necessary to determine the functional significance of these associations.

In summary, we have empirically validated an approach for the specific detection of ACE2 protein at the surface of single cells, revealing a surprising heterogeneity in cellular ACE2 expression within HuH7 and Calu-3 cells. Characterization of the subpopulation of ACE2-expressing cells supported an epigenetic mechanism of regulation, with RNA-seq profiling identifying broad transcriptome changes correlated with ACE2 expression. These findings advance our molecular understanding of ACE2 expression and should serve as a valuable resource for future studies of *ACE2* gene regulation.

## MATERIALS AND METHODS

### Cell culture and reagents

HuH7, Caco-2, Calu-3, HuH7.5.1, VeroE6, HepG2, and HEK293T cells were cultured in DMEM supplemented with 10% fetal bovine serum (FBS) and penicillin/streptomycin (Thermo Fisher Scientific, Waltham MA). CRISPR constructs were generated by cloning an *ACE2*-targeting gRNA sequence [TACCAAGCAAATGAGCAGGG] or a nontargeting control gRNA sequence [CGTGTGTGGGTAAACGGAAA] into Esp3I (New England Biolabs, Ipswich MA) sites of the pLentiCRISPRv2 backbone (Addgene, Watertown MA, #52961, gifted by Feng Zhang). An *ACE2* overexpression construct was generated by HiFi DNA assembly (New England Biolabs #E2621) of the human *ACE2* coding sequence (Sino Biological, Beijing China, #HG10108-M) into LeGO-iC2 (Addgene #27345, gifted by Boris Fehse) with simultaneous replacement of mCherry with a blasticidin resistance cassette. Lentivirus was generated and used to genetically engineer cell lines as previously described^57^. Primers used for qRT-PCR were: *ACE2*-fwd [5’ -AAACATACTGTGACCCCGCAT-3’], *ACE2*-rev [5’ -CCAAGCCTCAGCATATTGAACA-3’], *ACTB*-fwd [5’ - CCCTGGACTTCGAGCAAGAG-3’], *ACTB*-rev [5’ -ACTCCATGCCCAGGAAGGAA].

### Immunoblotting

Cells were lysed in RIPA buffer (Thermo Fisher, *#*89900) supplemented with a protease inhibitor cocktail (Sigma-Aldrich, St. Louis MO, #11836170001). Protein lysates were resolved on 4-12% Bis-Tris gels, and transferred to nitrocellulose membranes. Washes were performed with TBS-T and blocking with 5% non-fat mlik in TBS-T. Primary antibodies were diluted in blocking buffer and incubated with membranes at 4°C overnight with agitation. For all main text figures, membranes were probed with primary antibodies against ACE2 (GeneTex, Irving CA, #GTX01160, 1:1000), GAPDH (Abcam, Cambridge UK, #181602, 1:5000), and β-actin (Santa Cruz Biotechnology, Dallas TX, #sc-47778, 1:5000). Additional ACE2 antibodies assessed by immunoblotting in this study (Figure S1) are: R&D Systems (Minneapolis MN, #AF933, 1:200), Santa Cruz (sc-73668, 1:500), Santa Cruz (sc-390851, 1:500), Abcam (#272690, 1:500), Abcam (#108252, 1:1000), R&D Systems (#MAB933, 1:250), R&D Systems (#MAB9332, 1:250), Sigma (#SAB3500978, 1:1000), Atlas Antibodies (Stockhold Sweden, #HPA000288, 1:500), ProteinTech (Rosemont IL, #21115-1-AP, 1:1000). All primary antibodies were diluted according to the manufacturer’ s recommendations when specified, and otherwise were diluted at 1:500. Secondary antibodies were diluted 1:3000 in blocking buffer and incubated at room temperature for 1 hr with agitation. Secondary antibodies were matched to the primary antibody and included HRP-conjugated goat anti-rabbit IgG (Bio-Rad Laboratories, Hercules CA, #1706515), goat anti-mouse IgG (Bio-Rad #1706516), and rabbit anti-goat IgG (Bio-Rad #1721034) Immunoblots were developed with SuperSignal West Pico PLUS Chemiluminescent Substrate (Thermo Fisher #34579) and visualized on a ChemiDoc MP instrument (Bio-Rad) with exposure times automatically selected to optimize band intensity relative to background.

### Flow cytometry

For flow cytometry experiments, cells were harvested with TrypLE Express (Thermo Fisher), centrifuged at 500x*g* for 5 min, washed with FACS buffer (PBS supplemented with 2% FBS), centrifuged again, and resuspended in FACS buffer at a density of ∼10-20 million cells/mL. Where indicated, cell permeabilization was performed by incubating cells in ice cold methanol for 10 min. All other experiments were performed on intact cells. Blocking was performed by incubating cells on ice in FACS buffer for 10 mins, then primary antibody was added at the indicated dilution and cells were incubated on ice for ∼45 mins with intermittent gentle agitation to keep cells in suspension. Cells were washed twice with excess FACS buffer, and incubated with secondary antibody diluted in FACS buffer for 30 mins on ice. Cells were washed with excess FACS buffer, centrifuged, and resuspended in FACS buffer containing SytoxBlue (Thermo Fisher #S34857) live/dead cell stain at 1:1000 dilution. After a 3-5 min incubation, cells were again washed with excess FACS buffer, centrifuged, resuspended in FACS buffer at a density of ∼10-20 million cells/mL and filtered into FACS tubes. Flow cytometry was performed on a BD FACSAria III, BioRad Ze5, or BD Fortessa instruments, with all data gated and analyzed using FlowJo v10 software. Cell sorting experiments were performed on the BD FACSAria III instrument. For single antibody FACS experiments, R&D Systems #MAB9332 was used at 1:50 dilution. FACS experiments comparing a panel of ACE2 antibodies were performed with: GeneTex #GTX01160, R&D Systems #AF933, Santa Cruz sc-73668, Santa Cruz sc-390851, Abcam #272690, Abcam #108252, R&D Systems #MAB933, Sigma #SAB3500978, Sigma Prestige #HPA000288, ProteinTech #21115-1-AP, Novus Biologicals #NBP2-72117C, and Cell Signaling #4355S. Secondary antibodies were matched to the corresponding primary antibodies and included: AlexaFluor647 goat anti-mouse IgG antibody (Fisher #A32728), AlexaFluor647 goat anti-rabbit IgG (Fisher #A32733), and AlexaFluor647 donkey anti-goat IgG (Fisher #A32849). For cell cycle analysis, cells were incubated for 10 min on ice with Propidium Iodide Cell Staining Solution (Cell Signaling #4087) prior to flow cytometry.

### Microscopy

Cells were seeded in 35 mm poly-D lysine-coated glass bottom dishes (MatTek, Ashland MA, P35GC-1.5-14-C) so that they were ∼70-90% confluent on the day of staining. Cell monolayers were briefly washed with 4°C PBS then fixed with 4% paraformaldehyde in PBS for 10 min at room temperature and washed twice with PBS. All subsequent steps were performed at room temperature. For experiments involving intracellular staining, cells were then incubated with 0.1% saponin (Sigma-Aldrich) in PBS for 5 min and washed twice with PBS. Blocking was performed with 2% FBS in PBS for 15 min. Blocking buffer was aspirated and cells were incubated for 45 min with primary antibody (ACE2 R&D #MAB9332 and/or GM130 Abcam #ab52649, both at 1:100 dilution in blocking buffer). Cells were washed three times with PBS and incubated for 30 min with secondary antibody (AlexaFluor488-conjugated goat anti-mouse Thermo Fisher #A32723 and/or AlexaFluor594-conjugated donkey anti-rabbit Thermo Fisher #A32754 both at 1:500 dilutions in blocking buffer). Cells were washed three times with PBS and incubated with either DAPI (Thermo Fisher #62248) 1 µg/mL in PBS for intracellular staining or Hoechst 33342 (Thermo Fisher #H3570) at 1 µg/mL in PBS for nonpermeabilized cells. Images were obtained on a Nikon A1R confocal microscope and analyzed with Nikon NIS-Elements software.

### ACE2 activity assay

ACE2 enzymatic activity was measured using a fluorometric assay kit (BioVision, Milpitas CA, #K897). Cells were freshly harvested and ACE2 activity was measured in triplicate according to manufacturer protocol. Fluorescence data was collected on a ThermoMax microplate reader (Molecular Devices, San Jose CA).

### RNA analysis

Cellular total RNA was prepared from 2-4×10^6^ cells for each sample using the RNeasy Plus Micro kit (Qiagen, Hilden Germany, #74034). For qRT-pCR, cDNA was prepared using the SuperScript III first-strand synthesis kit (Thermo Fisher, #18080051), amplified with indicated primers using the *Power* SYBR Green PCR Master Mix (Thermo Fisher, # 4367659), and analyzed by QuantStudio 5 Real-Time PCR (Thermo Fisher). For RNA-seq analysis, RNA quality was assessed by electrophoresis on a TapeStation instrument (Agilent, Santa Clara CA). Samples with RNA Integrity Numbers of 8 or greater were prepared for sequencing. Polyadenylated mRNA was purified from 450 ng of total RNA with the NEBNext Poly(A) mRNA Magnetic Isolation Module (NEB, #E7490). Sequencing libraries were prepared with random primers using the NEBNext Ultra II Directional RNA Library Prep Kit for Illumina (NEB, #E7760L) with NEBNext Multiplex Oligos (NEB, #E6440L). Final libraries were analyzed by TapeStation electrophoresis and KAPA library quantification (Roche, Basel Switzerland, KK4835). The samples were pooled and sequenced on a NovaSeq instrument (Illumina, San Diego CA) S4 flow cell with 75 bp single end reads. Individual reads were aligned to the human genome version GRCh38.p13 with STAR version 2.7.1a^58^. Read counts were generated by HTSeq_count version 0.9.1^59^ and TPM values by Salmon version 1.4.0^60^. Differential gene expression analysis was performed using DESeq2^61^. For *ACE2* exon usage analysis, individual reads were aligned to the *ACE2* NCBI Refseq genomic sequence NG_012575.2 using TopHat version 2.1.1^62^. Isoform expression was visualized with the Sashimi utility of the MISO package version 0.5.4^63^.

### Gene set enrichment analysis

Gene set enrichment analysis of RNA-seq data was performed using GSEA v4.1.0^33^. Gene sets were chosen from the Molecular Signatures Database (MSigDB, v7.2), and included the Hallmark gene set^34^ for pathway analysis and the Gene Transcription Regulation Database (GTRD) gene set^35^ for transcription factor analysis. For all analyses, settings included 1000 gene set permutations, weighted enrichment with the Signal2Noise metric, and gene set sizes between 15 and 500 genes. An FDR cutoff of 25% was used to select significant gene sets, which were then ranked based on a normalized enrichment score.

## AUTHOR CONTRIBUTIONS

E.J.S. and B.T.E. conceived of the study, collected and analyzed data, and prepared the manuscript.

## ADDITIONAL INFORMATION

This research was supported by NIH grant K08-HL148552 (BTE). The authors declare no competing interests.

## ACKNOWLEDGMENTS

We thank Dave Siemieniak for assistance with the bioinformatics analysis described in this manuscript.

---

**Supplemental Figure 1.**
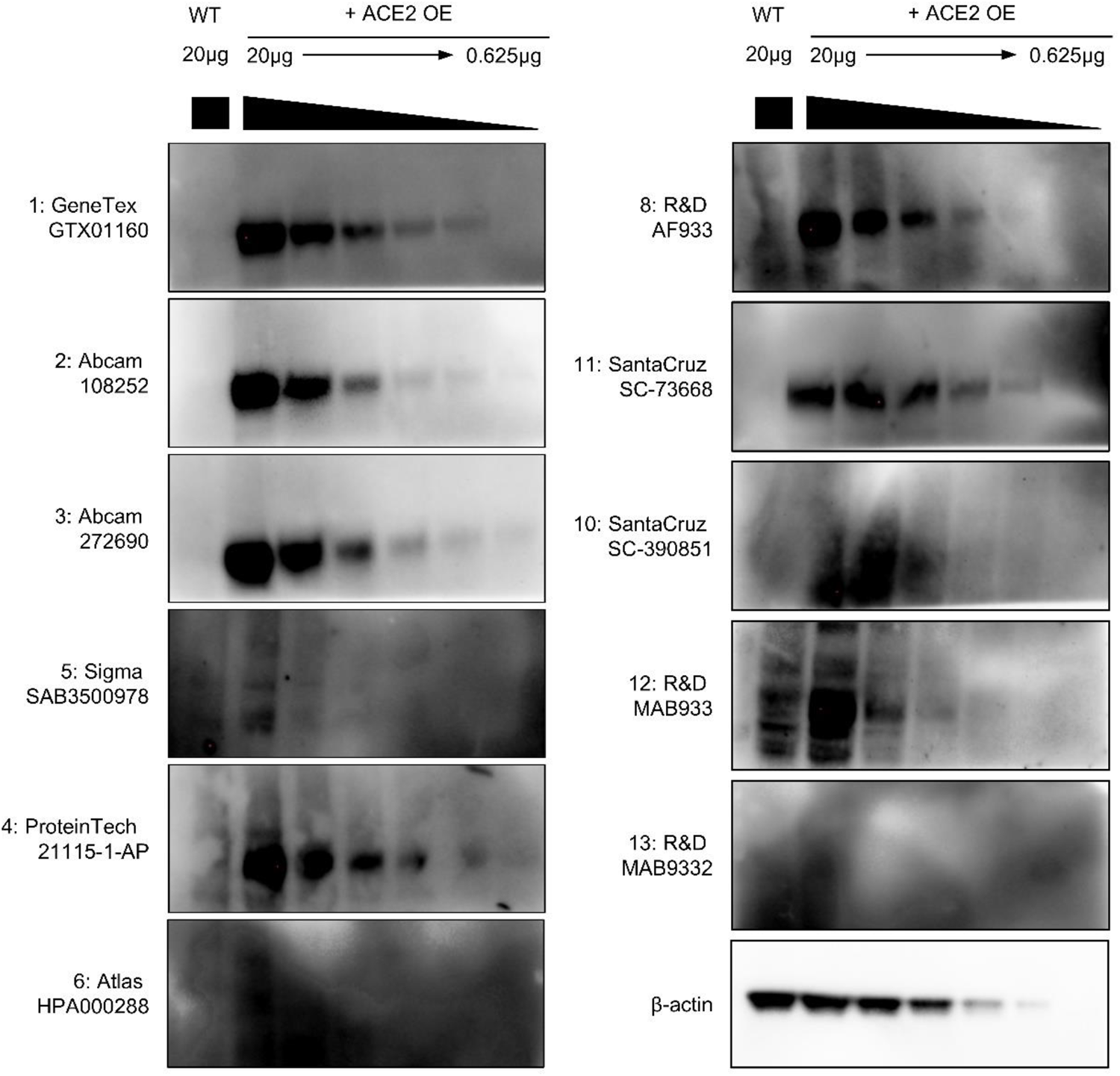
Assessment of ACE2 antibody sensitivity and specificity by immunoblotting. Representative immunoblots for ACE2 using each of 11 different commercial antibodies, comparing 20 µg of lysate of HEK293T parental cells to a dilution gradient of 20 µg to 0.625µg of lysates from ACE2-overexpressing HEK293T cells.

**Supplemental Figure 2.**
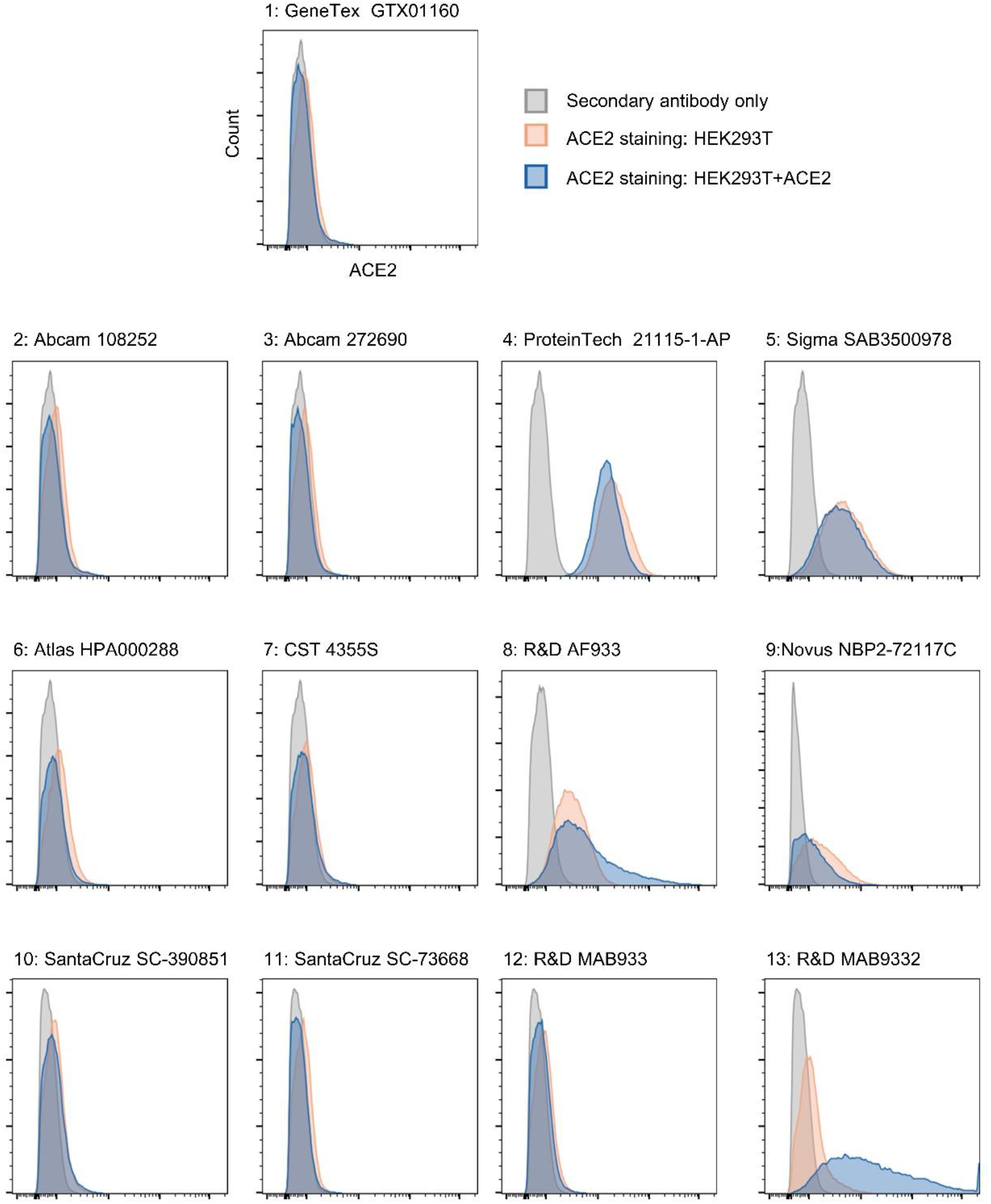
Assessment of antibody specificity for detection of ACE2 by flow cytometry. Representative histograms using each of 13 different commercial ACE2 antibodies (summarized in Figure 1C and Table 1), comparing ACE2 staining of ACE2-overexpressing HEK293T cells to ACE2 staining and secondary antibody only staining of parental cells.

**Supplemental Figure 3.**
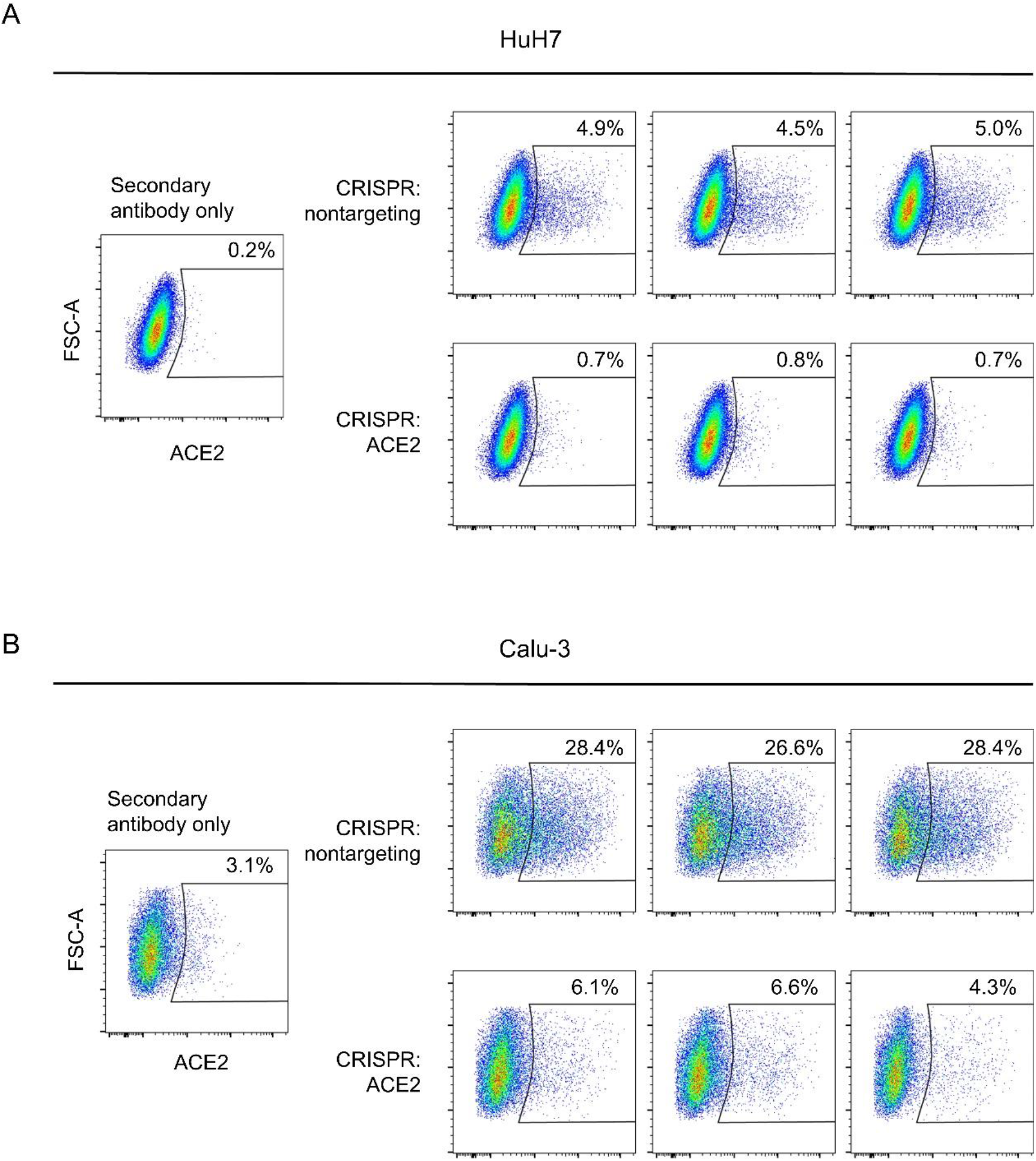
Quantification of ACE2-positivity in HuH7 and Calu-3 cells by flow cytometry. Individual plots for three biological replicates of ACE2 staining by flow cytometry in HuH7 and Calu-3 cell lines generated by CRISPR with either a nontargeting guide (NT) or *ACE2*-targeting gRNA, (summarized in Figure 2C).

**Supplemental Table 1. Transcriptomes of ACE2-negative and ACE2-positive HuH7 cells**. Mapped reads for each transcript identified by RNA-seq of cells collected by flow cytometry sorting based on ACE2 surface abundance for each of 4 independent biologic replicates.

**Supplemental Table 2. Transcriptomes of parental wild-type and serially ACE2-enriched HuH7 cells**. Mapped reads for each transcript identified by RNA-seq of parental HuH7 cells and cells harvested after 3 rounds of serial ACE2 enrichment by flow cytometry for each of 3 independent biologic replicates.

**Supplemental Table 3. Differential gene expression analysis of sorted HuH7 ACE2-negative and ACE2-positive cells**. DESeq2 output of differentially expressed genes among RNA-seq profiles of cells collected by flow cytometry sorting based on ACE2 surface abundance for each of 4 independent biologic replicates.

**Supplemental Table 4. Differential gene expression analysis of HuH7 parental wild-type and serially ACE2-enriched cells**. DESeq2 output of differentially expressed genes among RNA-seq profiles of parental HuH7 cells and cells harvested after 3 rounds of serial ACE2 enrichment by flow cytometry for each of 3 independent biologic replicates.

**Supplemental Table 5. Hallmark gene set analysis of ACE2-correlated transcripts**.

GSEA output with normalized enrichment score (NES), nominal p-value, and FDR for hallmark gene set analysis of either singly sorted ACE2-negative and ACE2-positive HuH7 cells, or wild-type and serially sorted ACE2-enriched HuH7 cells. Positive NES values indicate enrichment and negative values depletion in ACE2-expressing cells.

**Supplemental Table 6. Transcription factor target analysis of ACE2-correlated transcripts**. GSEA output with normalized enrichment score (NES), nominal p-value, and FDR for GTRD gene set analysis of either singly sorted ACE2-negative and ACE2-positive HuH7 cells, or wild-type and serially sorted ACE2-enriched HuH7 cells. Positive NES values indicate enrichment and negative values depletion in ACE2-expressing cells.

